# Intra-nanoparticle Drug-protein Interactions Mediate Sequential Therapeutic Release

**DOI:** 10.64898/2026.06.12.731961

**Authors:** Huibin Chang, Anup Dey, Tengfei Ma, Ingrid Oprea, Rachel Y. Zhang, Sufeng Zhang

**Author notes:** To whom correspondence should be addressed: **Corresponding Author Sufeng Zhang** - Department of Biomedical Engineering, Stony Brook University, Stony Brook, 11794, USA; Department of Pharmaceutical Sciences, Stony Brook University, Stony Brook, 11794, USA; Division of Gastroenterology, Hepatology and Endoscopy, Department of Medicine, Brigham and Women’s Hospital, Boston, MA 02115, USA, Harvard Medical School, Boston, MA 02115, USA.

## Abstract

Controlled drug delivery systems have important implications in many therapeutic applications to improve patient health, but it remains challenging to deliver therapeutics to specific target sites while being released at controlled rates to minimize the off-target accumulation. Here, we report a drug (e.g., vancomycin)–modulating other drug release strategies in human serum albumin (HSA)-based nanoparticles (NPs) to achieve multiple drug encapsulation and controlled release of drugs. Without external stimuli but only with water, the release kinetics of drugs such as sulfasalazine and epidermal growth factor can be modulated by adding vancomycin. The mechanistic study suggests that the release kinetics are related to the secondary structure of albumin through the interactions between drugs and HSA. Our strategy achieved controlled drug delivery through modulating the secondary structure of albumin with a water-soluble drug as a modulator, which provides a promising way to simultaneously deliver multiple drugs while releasing in a predicted and sustained manner.

## Introduction

Inflammatory bowel disease (IBD) is a chronic inflammatory disease of the gastrointestinal tract and consists of two main types: Crohn’s disease (CD) and ulcerative colitis (UC)^1^. For patients with CD, discontinuous inflamed spots are usually observed in the digestive tract, ranging from the mouth to the anus, while with UC, a continuous inflamed area only exists in the large intestine^2^. In the past century, the prevalence of IBD was predominant in developed countries, but has been rising in newly industrialized countries^3, 4^. Although the exact etiologies remain uncertain, the pathogenesis of IBD likely involves genetic, environmental, microbial, and immunological factors^5-7^. Current drug treatments aim not only to induce and maintain patients in remission and ameliorate the disease’s secondary effects but also to potentially achieve mucosal healing^8, 9^. Therefore, delivering specific therapeutics to the target site at a controlled dose with minimal off-target toxicities could potentially improve the health condition of many patients^10^.

The current frontline management of IBD relies largely on monotherapies, including 5-aminosalicylates (5-ASAs), corticosteroids, immunomodulators, and biologics targeting individual inflammatory mediators such as tumor necrosis factor-alpha (TNF-α)^11, 12^. Although these agents can induce clinical responses, durable remission remains difficult to achieve, in part because IBD arises from multiple interconnected pathogenic processes that cannot be fully addressed through inhibition of a single target^13-17^. This recognition has driven growing interest in combination therapeutic strategies that simultaneously modulate different disease pathways to achieve more comprehensive disease control^18^. Sulfasalazine (SSZ) was among the first effective pharmacological treatments for IBD and remains an important therapeutic option, particularly for patients with mild-to-moderate UC^19, 20^. SSZ is made up of two components: 5-ASA (the active therapeutic moiety) and sulfapyridine^5^. Although it is not active therapeutically, overdose of sulfapyridine may cause severe side effects. Vancomycin (VAN) is a glycopeptide antibiotic and is used to treat severe infections caused by Gram-positive bacteria^21^. Previous studies showed that patients achieved clinical remission and mucosal healing with VAN; however, 500 mg of VAN was administered twice a day orally^22^. Recombinant peptides are increasingly being used for clinical purposes. For example, epidermal growth factor (EGF) is an effective treatment for patients with active left-sided UC when they receive daily enemas of 5 µg of EGF in 100 ml of an inert carrier for 14 days^13^. To preserve the therapeutic effect of drugs without the side effects, controlled release formulations with minimal drug loading are necessary in the small intestine and colon^5^.

NP-based drug delivery systems represent a powerful platform for improving the therapeutic efficacy of a wide range of diseases by enhancing drug stability, prolonging circulation time, increasing bioavailability, and enabling targeted delivery^23, 24^. Among various nanocarriers, human serum albumin (HSA) NPs have attracted significant attention owing to their excellent biocompatibility, biodegradability, non-immunogenicity, and intrinsic drug-binding capability^25, 26^. HSA NPs can efficiently encapsulate both hydrophobic and hydrophilic therapeutics, protect them from premature degradation, and provide sustained and controlled drug release. These advantages are particularly beneficial for IBD, where conventional therapies often suffer from drug retention at non-inflamed intestinal sites, thereby limiting therapeutic efficacy and leading to severe systemic side effects. Furthermore, HSA NPs were reported to enhance drug accumulation at inflamed intestinal tissues and improve local therapeutic retention^27^, which could potentially reduce systemic exposure and maximize therapeutic outcomes. Consequently, HSA-based nanoplatforms provide a promising platform to enable co-delivery of multiple therapeutics while improving drug accumulation and retention in inflamed intestinal tissues.

In this work, we report HSA-based NPs capable of modulating triple-drug delivery through intra-NP drug interactions. Specifically, VAN was found to actively regulate the release kinetics of encapsulated SSZ and EGF, enabling sequential and controlled therapeutic release without external stimuli. Using water as the triggering factor, the release behavior of therapeutics was investigated through *in vitro* experiments and computational modelling. Our goal is to establish a conceptually innovative strategy for modulating drug release in nanomedicine that could be broadly applicable to the design of novel combination therapies.

## Results and Discussion

Three different drugs, including small-molecule drugs (VAN and SSZ) and proteins (EGF), were studied for the loading and releasing properties of single-, dual-, and triple-drug loaded HSA NPs **(Figure 1)**. HSA NPs were prepared through three main steps: reduction of HSA intramolecular disulfide bonds using glutathione (GSH), formation of HSA NPs via ethanol desolvation, and removal of excess GSH and ethanol through dialysis ^28^. After optimization, HSA at a concentration of 80 mg/mL in water was chosen based on morphology and size measurement. The representative morphology of HSA NPs showed a uniform spherical shape **(Figure S1 a, b)**. In an aqueous solution, the hydrodynamic size of HSA NPs is ∼150 nm with a low polydispersity index (PDI, ∼0.05 in **Figure S1 c, d**). HSA NPs exhibited a positive surface charge with low variability in zeta potential measurements (**Figure S1 e, f**), indicating the synthesized NPs are relatively stable in water. The positive zeta potential of the HSA NPs is likely associated with GSH-mediated disulfide bond reduction during NP synthesis^29^.

**Figure 1.**
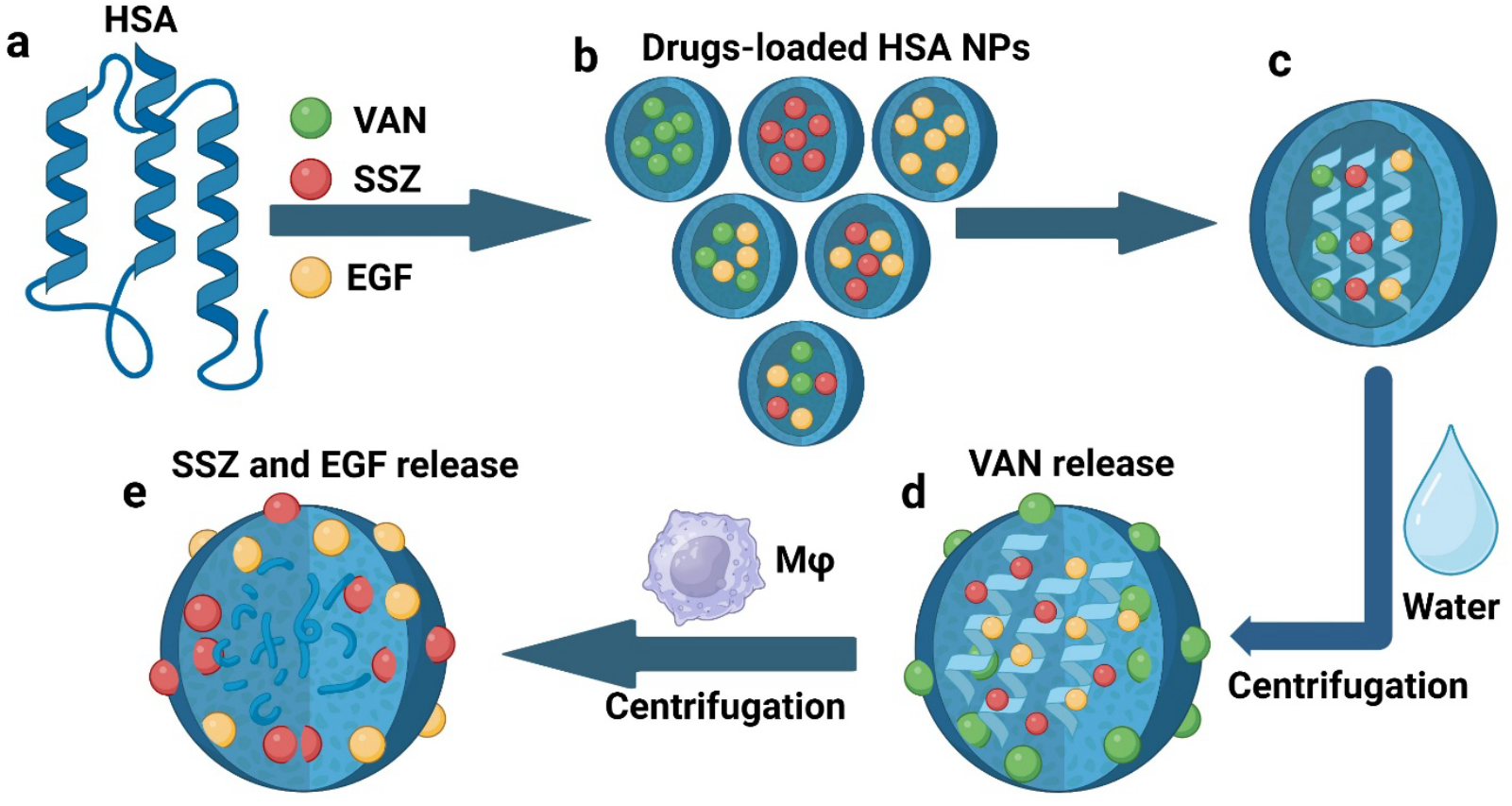
Schematic illustration of drug-loaded HSA NPs with sequential drug release. (a) Human serum albumin (HSA) was used to synthesize NPs for three different drugs. (b) Illustration of single, dual, and triple-drug loaded HSA NPs. (c - e) Centrifugation in water quickly released the water-soluble drug, VAN, and biological factors from the supernatant of activated macrophages stimulated the release of SSZ and EGF from HSA NPs.

First, drugs (VAN, SSZ, and EGF) were individually loaded into HSA NPs to check the loading and release efficiency of each drug from NPs. The drug-loaded HSA NPs were spherical in shape **(Figure 2a-c)** and showed an increase in size when VAN, SSZ, or EGF was loaded into NPs **(Figure 2d and Figure S2)**, ranging from ∼130 nm to ∼230 nm. The zeta potential of the drug-loaded NPs showed a similar positive charge as compared to HSA NPs **(Figure 2e)**. These results suggest that HSA can effectively encapsulate diverse therapeutic cargos without significantly altering the surface properties, demonstrating its potential as a versatile drug carrier. To determine the encapsulation efficiency (EE) of each drug, HSA NPs were digested by GSH, and the amount of drug was quantified by high-performance liquid chromatography (HPLC) for VAN and SSZ, and by enzyme-linked immunosorbent assay (ELISA) for EGF. For single-drug loaded NPs, the result showed an EE of 8.1% ± 2.1%, 70.4% ± 4.4%, and 16.2% ± 2.0% for VAN, SSZ, and EGF, respectively **(Figure 2f)**. *In vitro* drug release was investigated in the presence of RAW264.7 macrophages (MΦ) with lipopolysaccharide (LPS) activation to simulate the inflammatory condition^27^. The results show that a higher amount of LPS-activated MΦ supernatant led to higher amount of drug release **(Figure 2g-i)**, but the release percentage of VAN was much higher than that of SSZ. Surprisingly, unlike SSZ and EGF showing less release in water, VAN was released closer to 100% from the VAN-loaded NPs in water after centrifugation **(Figure 2i)**.

**Figure 2.**
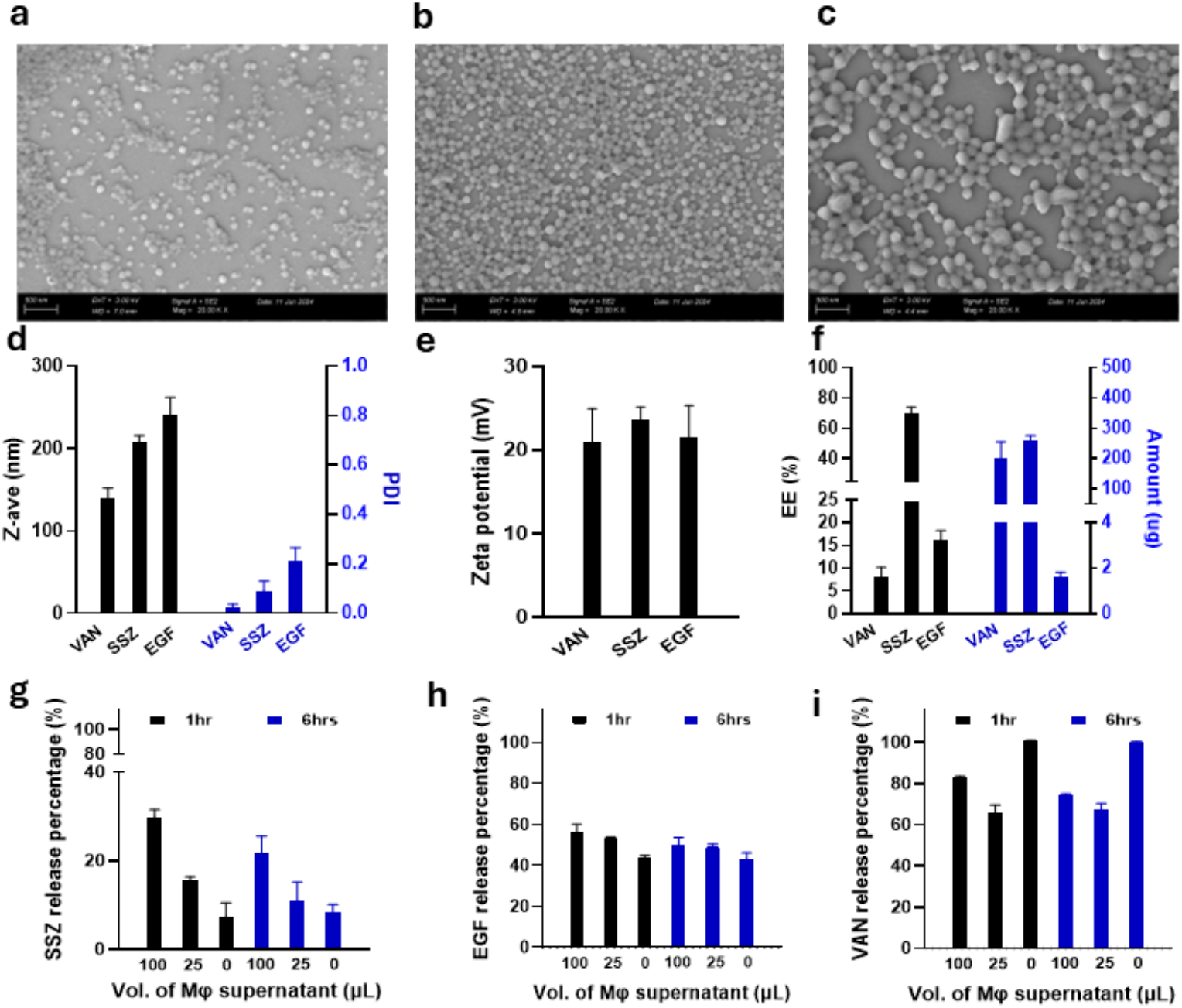
Characterization of single drug (VAN, SSZ, or EGF) loaded HSA NPs. (a-c) Representative images of (a) VAN-loaded HSA NPs, (b) SSZ-loaded HSA NPs, and (c) EGF-loaded HSA NPs. HSA concentration was 80 mg/mL in the NP formulations. (d-e) Hydrodynamic size and zeta potential of single-drug loaded HSA NPs (n ≧ 4). (f) EE of single-drug loaded HSA NPs (n=2). (g-i) *In vitro* drug release from single-drug loaded NPs in the presence of varying amounts of LPS-activated macrophage supernatant (n=2). Results are presented as mean ± SD.

Next, we chose to load drug combinations of (VAN+EGF) and (SSZ+EGF) as a combination of small-molecule drugs and biologics in NPs, which could potentially be used as a system to study the synergetic effects of drugs. The amount of each drug (VAN, SSZ, and EGF) in the dual-drug loaded NPs was determined based on the optimized EE of the corresponding single-drug loaded NPs. The dual-drug loaded NPs showed spherical shapes, and the (SSZ+EGF)-loaded NPs exhibited larger diameter than the (VAN+EGF)-loaded NPs **(Figure 3a, b)**, which was in accordance with the hydrodynamic size measurements: about 280 nm for (SSZ+EGF)-loaded NPs and about 180 nm for (VAN+EGF)-loaded NPs **(Figure 3c and Figure S3)**. This difference in size was probably due to the different physicochemical properties of the two drugs: SSZ is a more hydrophobic molecule than VAN, interacting strongly with the hydrophobic pocket of HSA NPs^30^. This stronger interaction formed a rigid molecular complex, which can enhance the size of NPs. While VAN is more hydrophilic in nature, this resulted in more compact and smaller NPs than (SSZ+EGF)-loaded NPs. The dual-drug-loaded NPs exhibited a zeta potential similar to that of single-drug-loaded NPs (∼20 mV) **(Figure 3d)**. For the (VAN+EGF) loaded NPs, the EE was 8.1% ± 1.5% for VAN and 9.8% ± 2.0% for EGF, respectively **(Figure 3e)**. For the (SSZ+EGF) loaded NPs, the EE was 56.6% ± 1.9% for SSZ and 46.4% ± 10.8% for EGF, respectively **(Figure 3f)**.

**Figure 3.**
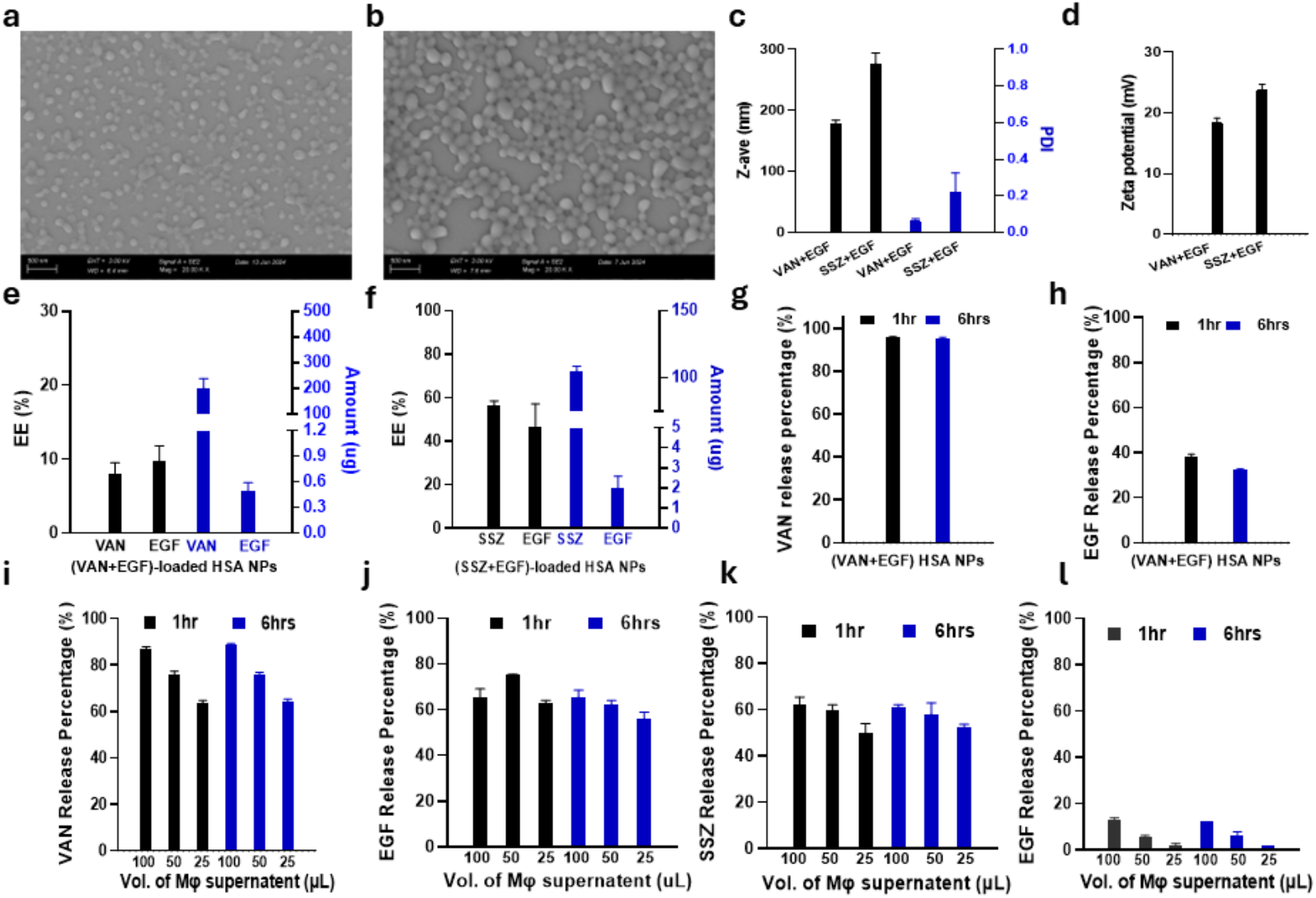
Characterization of dual drugs (VAN+EGF and SSZ +EGF) loaded HSA NPs. (a, b) Representative SEM images of (a) (VAN+EGF)-loaded HSA NPs and (b) (SSZ+EGF)-loaded HSA NPs. The HSA concentration was 80 mg/mL in the NP formulations. (c, d) Hydrodynamic size and zeta potential of the dual-drug loaded HSA NPs (n = 4). (e, f) EE determination of the dual-drug loaded HSA NPs (n = 4). (g, h) *In vitro* drug release of VAN and EGF from (VAN+EGF)-loaded HSA NPs in the presence of water (n = 4). (i-l) *In vitro* drug release from dual-drug loaded NPs in the presence of varying amounts of LPS-activated macrophage supernatant. (i) VAN from (VAN+EGF)-loaded HSA NPs, (j) EGF from (VAN+EGF)-loaded HSA NPs, (k) SSZ from (SSZ+EGF)-loaded HSA NPs, and (l) EGF from (SSZ+EGF)-loaded HSA NPs (n = 2). Results are presented as mean ± SD.

Since water led to the release of VAN from single-drug loaded NPs after centrifugation, (VAN+EGF)-loaded NPs were first investigated for drug release in the presence of water. The results showed that closer to 100% VAN was released from NPs after 1 hour incubation at 37°C with 100 rpm shaking (**Figure 3g**). Due to its hydrophilic nature, VAN does not readily partition into the inner hydrophobic cavities of HSA NPs. Instead, it is likely localized within the hydrophilic core or associated with the NP surface through electrostatic interactions. Then, the drug release was further investigated in the presence of supernatant from LPS-activated MΦ **(Figure 3i-l)**. For (VAN+EGF)-loaded NPs, compared with water, LPS-activated MΦ supernatant did not accelerate VAN release and produced lower overall VAN release percentage, despite a dose-dependent increase in VAN release with increasing amounts of LPS-activated MΦ supernatant **(Figure 3i)**. For (SSZ+EGF)-loaded NPs, a higher amount of LPS-activated MΦ supernatant led to higher SSZ release; however, the release percentage was lower than that of VAN **(Figure 3k)**. Due to the difference in release kinetics between VAN and SSZ, the EGF release from the dual-drug loaded NPs was significantly affected: the release percentage of EGF from (VAN+EGF)-loaded NPs was much higher than that of (SSZ+EGF)-loaded NPs (**Figure 3j, l**).

To understand how VAN modulated the release of other drugs from HSA NPs, we investigated the underlying mechanism. First, we investigated the effect of VAN on HSA conformational changes during NP synthesis using GSH as the reducing agent by circular dichroism (CD) spectroscopy, a technique highly sensitive to protein secondary structure and widely used to study drug–albumin interactions^31^. The CD spectra showed characteristic peaks at 222 nm and 208 nm, which are commonly used to assess α-helix content and correspond to n-π* and π-π* transitions, respectively. Compared with pure HSA, HSA treated with GSH or GSH+VAN exhibited an upward shift in the CD spectra, with reduced negative ellipticity at 208 and 222 nm (**Figure 4a, b)**, suggesting decreased α-helical content and consequent destabilization of the HSA structure. In addition, the concentrations of SSZ and VAN also influenced the α-helical content of HSA **(Figure S4)**. Second, Fourier-transform infrared spectroscopy (FTIR) was used to investigate drug-HSA interactions. The amide I (∼1,600-1700 cm^−1^) and amide II (1,548 cm^−1^) bands provide information on the secondary structure of HSA^32^, with amide I being particularly sensitive to protein conformational changes. Both VAN- and SSZ-loaded HSA exhibited shifts in the amide I peak (∼ 1653 cm^−1^), indicating alterations in the HSA secondary structure following drug incorporation (**Figure 4c**). Notably, HSA+SSZ showed a larger amide shift than HSA+VAN, suggesting stronger interactions between SSZ and HSA. Protein folding and structural stability during NP synthesis are governed by intra- and intermolecular interactions, including hydrogen bonding, hydrophobic, polar, and van der Waals interactions^33, 34^. Therefore, while VAN and SSZ may induce comparable secondary structural changes in HSA, the smaller amide I shift associated with VAN suggests weaker VAN–HSA interactions relative to SSZ–HSA. Third, we measured the hydrodynamic size and zeta potential of VAN-loaded HSA NPs following incubation at 37°C with shaking (100 rpm) for up to 60 min. The largely unchanged hydrodynamic size and zeta potential indicated that VAN release occurred without substantial alteration of NP morphology or surface characteristics (**Figure 4d, e**). To assess morphological changes associated with VAN release, NPs were characterized by SEM. Following incubation in water for 20 min at 37°C, some NPs displayed roughened surfaces, and smaller spherical particles were observed (**Figure 4f**), compared with the smooth morphology and relatively uniform size of VAN-loaded NPs before incubation (**Figure 2a**). Based on the hydrodynamic size measurements and SEM observations, we reasoned that disruption of the HSA secondary structure (α-helix) may facilitate water diffusion into the HSA NPs, thereby generating a VAN concentration gradient that could drive VAN diffusion from the NP interior into the surrounding water **(Figure 4g)**.

**Figure 4.**
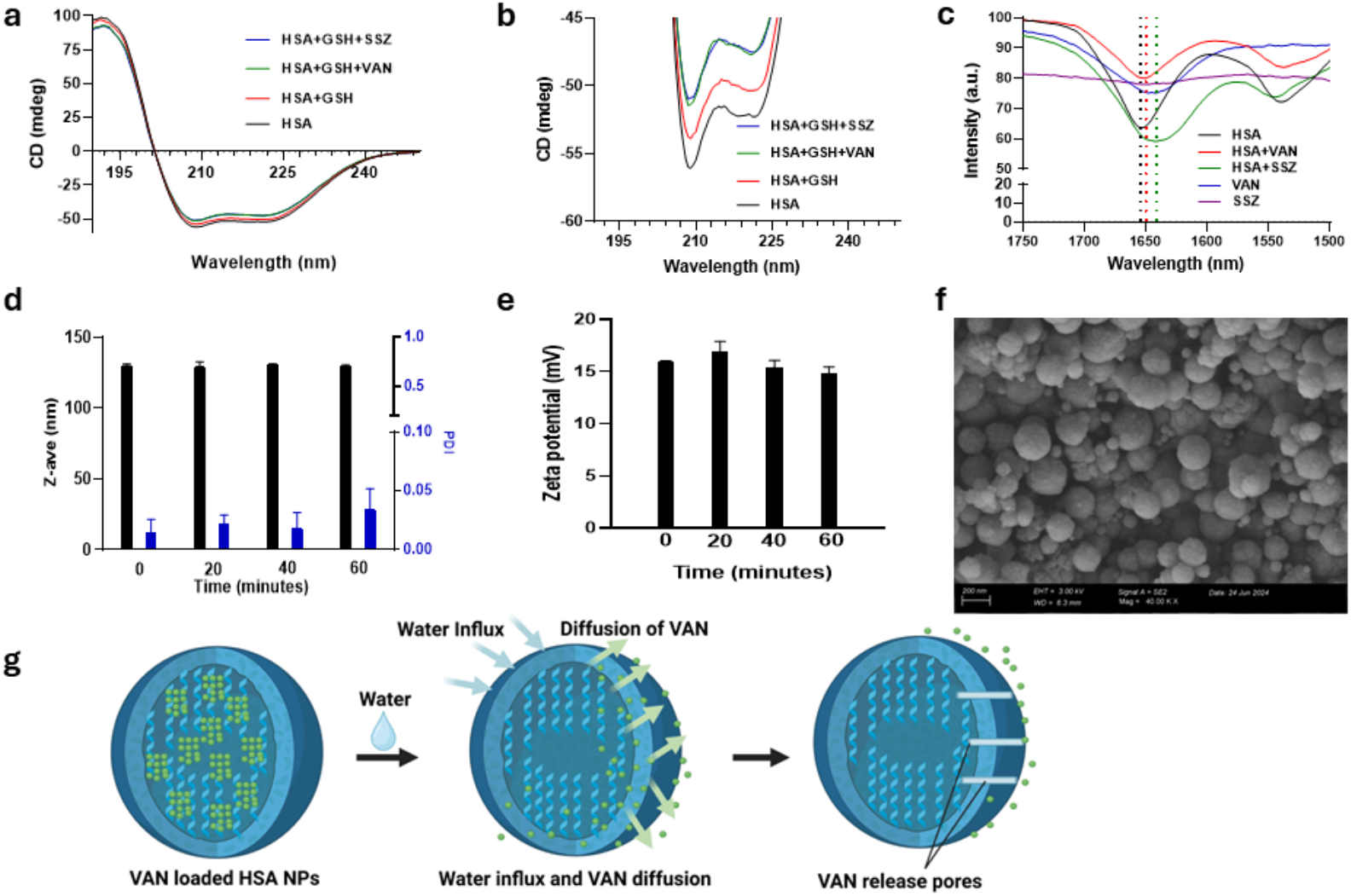
Mechanistic investigation of VAN release from HSA NPs. (a, b) Circular Dichroism (CD) Spectra of HSA, HSA+GSH, and HSA+GSH+VAN. (c) Fourier Transform Infrared (FTIR) spectra of HSA, VAN-loaded HSA, and SSZ-loaded HSA. (d, e) Hydrodynamic size and zeta potential of VAN-loaded HSA NPs in the presence of water (1:1 V/V%, 37°C) for 0, 20, 40, and 60 minutes. (f) Representative morphology of VAN-loaded HSA NPs in the presence of water for 20 minutes at 37°C. (g) Schematic of possible VAN-release mechanism from HSA NPs in the presence of water at 37°C.

We next synthesized (VAN+SSZ+EGF) triple-drug loaded NPs and evaluated the release of each drug, compared with (SSZ+EGF) dual-drug loaded NPs, to further investigate how VAN modulates the release of other drugs. The triple-drug loaded NPs exhibited a diameter of ∼ 200 nm, positive surface charges, and spherical morphology **(Figure 5 a-c** and **Figure S5)**, comparable to those of single- and dual-drug loaded NPs. In addition, incorporation of three drugs into HSA NPs did not substantially alter the EE of each drug compared with the corresponding single-drug loaded HSA NPs **(Figure 5d)**. To confirm the effect of VAN on modulating the release of other drugs from HSA NPs, *in vitro* release studies were performed using both (VAN+SSZ+EGF) triple-drug loaded NPs and (SSZ+EGF) dual-drug loaded NPs. The NPs were first incubated in water, followed by the addition of supernatant from LPS-activated macrophages **(Figure 5e)**. Compared with (SSZ+EGF)-loaded NPs, (VAN+SSZ+EGF)-loaded NPs showed higher release percentages of both SSZ and EGF, suggesting that early VAN release induces structural changes in HSA NPs that facilitate subsequent SSZ and EGF release in the presence of supernatant from LPS-activated macrophages **(Figure 5f-h)**.

**Figure 5.**
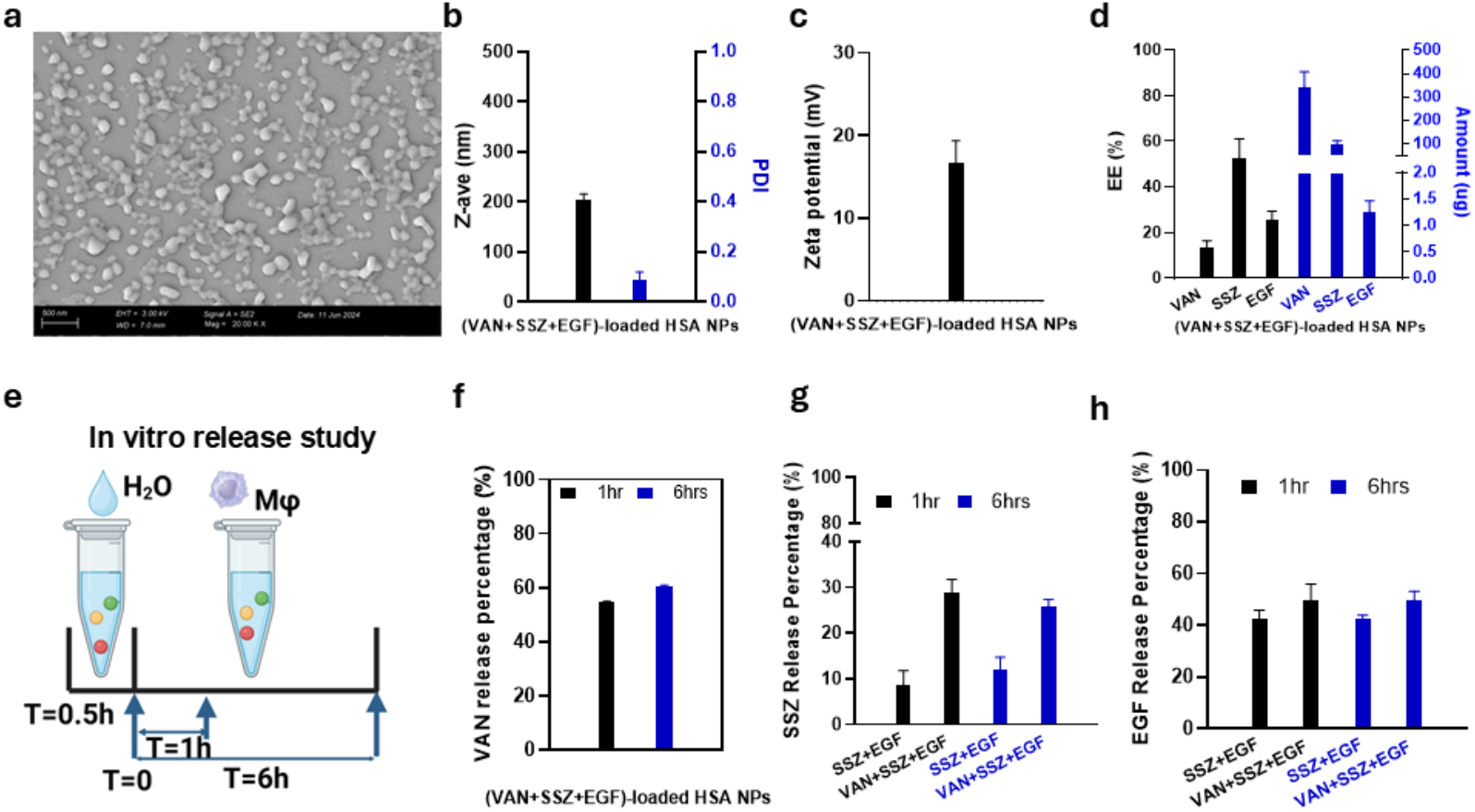
Characterization of triple-drug (VAN+SSZ+EGF) loaded HSA NPs with modulated drug release. (a) SEM image of the triple-drug loaded HSA NPs. (b, c) Hydrodynamic size and zeta potential measurements of the triple-drug loaded HSA NPs (n = 4). (d) EE of each drug for the triple-drug loaded HSA NPs (n = 4). (e) Schematic of *in vitro* drug release from HSA NPs. Drug-loaded HSA NPs were first incubated in water for 30 min, followed by the addition of supernatant from LPS-activated macrophages. (f-h) Release of VAN, SSZ, and EGF from the (VAN+SSZ+EGF)-loaded HSA NPs in comparison with (SSZ+EGF)-loaded HSA NPs (n = 4). Results are presented as mean ± SD.

To complement the spectroscopic evidence, we further compared the relative drug-HSA interactions by molecular docking, production molecular dynamics (PMD), and MM-GBSA binding free energy analysis. In the single-drug HSA model, the calculated GBSA binding energy for VAN and HSA was more favorable than that for SSZ and HSA over the 5001-10001 frame analysis window (-23.3 kcal mol^-1^ for VAN-HSA versus -15.0 kcal mol^-1^ for SSZ-HSA). This result appears different from the higher SSZ EE and slower SSZ release observed experimentally, and may also differ from literature trends measured for drug-albumin binding in dilute solution: -5.16 kcal mol^−1^ for VAN and -8.59 kcal mol^−1^ for SSZ^31, 35, 36^. Several factors can explain this difference. First, the single-load calculation estimates a local interaction free energy for one docked pose on solvated monomeric HSA, whereas the nanoparticle experiments measure overall drug retention from a concentrated, crosslinked or aggregated HSA matrix during GSH-induced structural rearrangement, water penetration, and diffusion through the particle. Second, literature binding constants are commonly obtained under defined solution conditions and can reflect multiple HSA binding sites, ligand protonation state, albumin fatty-acid occupancy, salt concentration, and entropic contributions that are not fully captured by a single MM/GBSA endpoint calculation. Third, VAN is large, highly polar, and water soluble; it can form favorable local electrostatic and hydrogen-bonding contacts with HSA in the simulation while still partitioning readily into water and promoting HSA secondary-structure disruption, both of which favor rapid release from NPs. Therefore, a more favorable calculated VAN-HSA single-site binding energy does not necessarily imply stronger nanoparticle retention. Consistent with this interpretation, the dual-drug load GBSA model better matched the release experiments: SSZ and HSA/EGF showed a more favorable binding energy than VAN and HSA/EGF (-16.8 kcal mol^-1^ versus -12.1 kcal mol^-1^), in agreement with the higher EE and lower release efficiency observed for (SSZ+EGF)-loaded HSA NPs compared with (VAN+EGF)-loaded HSA NPs. Thus, the single-load simulation should be interpreted as a site-level drug-HSA affinity comparison, while the dual-drug load model is more representative of the experimental co-loaded NP release behavior **(Figure 6** and **Table S1)**.

**Figure 6.**
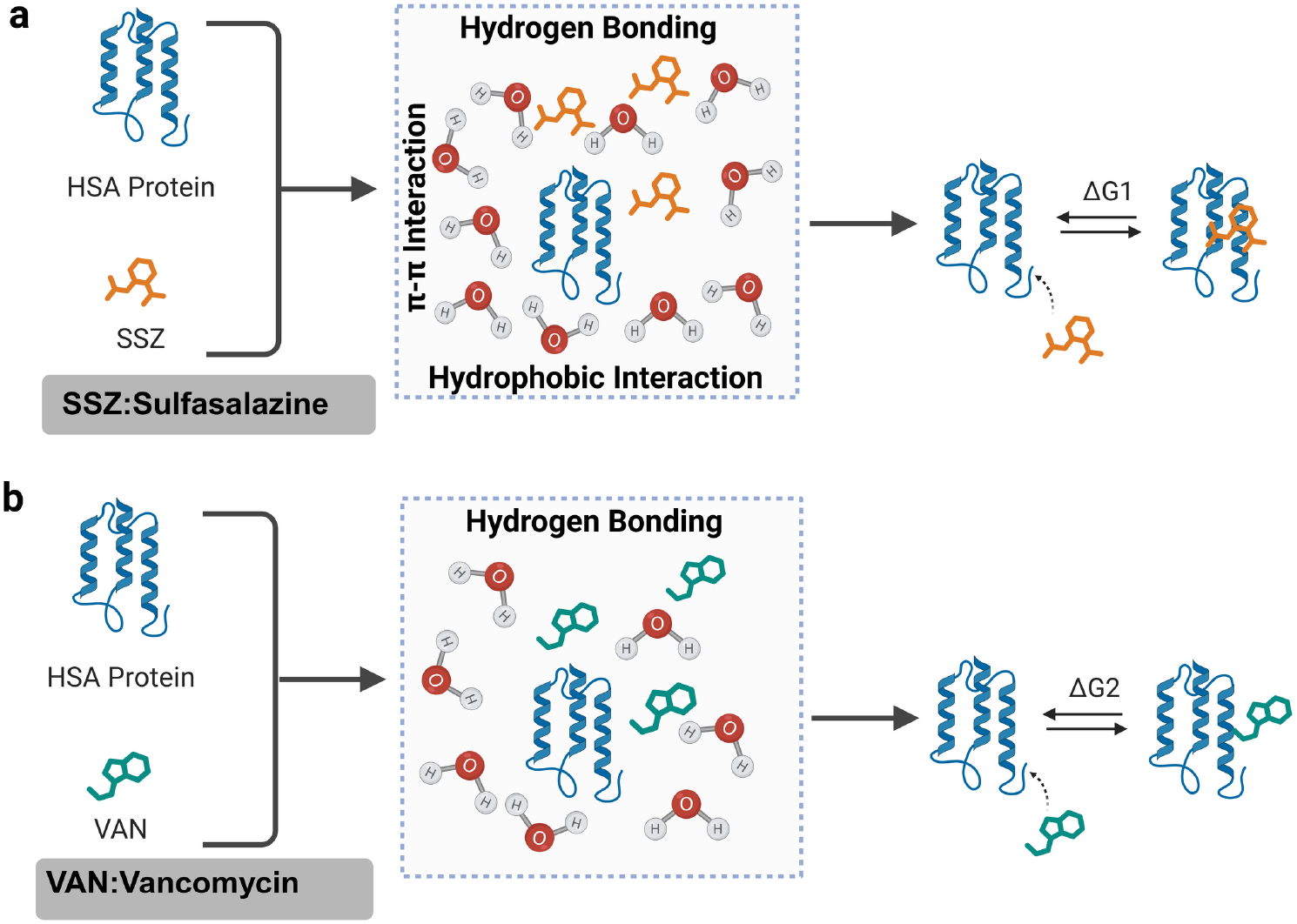
Schematic illustration of the differential interactions between SSZ and VAN with HSA based on molecular modeling analysis. (a) Interaction between SSZ and HSA NPs. (b) Interaction between VAN and HSA NPs. SSZ exhibited stronger interactions with HSA, potentially due to enhanced hydrogen-bonding and hydrophobic interactions within the albumin binding region. In contrast, VAN showed weaker interactions with HSA, which may contribute to its earlier release from HSA NPs and subsequent modulation of the release of other therapeutics (ΔG: binding free energy).

We also compared the triple-load model with the dual-load model to evaluate the effect of adding VAN into the SSZ+EGF-loaded system. In the 5001-10001 frame GBSA analysis, the calculated SSZ-HSA interaction became slightly less favorable in the triple-load model than in the dual SSZ+EGF model (-14.5 kcal mol^-1^ versus -16.8 kcal mol^-1^), while the calculated VAN-HSA interaction in the triple-load model was comparable to or slightly more favorable than that in the dual VAN+EGF model (-16.6 kcal mol^-1^ versus -12.1 kcal mol^-1^). This comparison suggests that the addition of VAN can weaken or perturb the SSZ-retaining HSA environment while maintaining favorable local VAN-HSA contacts, which is consistent with the experimental observation **(Figure 5f, g**) that triple-drug loading improves SSZ and EGF release compared with the SSZ+EGF dual-loaded NPs. Consequently, VAN release from multidrug-loaded HSA NPs may modulate the release behavior of co-encapsulated drugs, including SSZ and EGF.

## Conclusions

In summary, we developed a conceptually new controlled drug delivery system in which water-driven early VAN release modulates subsequent drug release from HSA-based NPs. We demonstrated that HSA NPs can simultaneously load versatile therapeutics, including small-molecule drugs and biologics (e.g., VAN, SSZ, and EGF). Water-driven VAN release was observed in both single- and dual-drug-loaded HSA NPs and was associated with VAN-mediated destabilization of the HSA secondary structure. Furthermore, we showed that water-driven VAN release in triple-drug-loaded HSA NPs can regulate the release of other drugs. Finally, molecular modeling, together with experimental studies, was used to probe the mechanism underlying this drug-mediated sequential release, suggesting that early VAN release can modulate the release behavior of subsequently released therapeutics. These findings highlight this drug-modulating drug release platform as a promising strategy for controlled drug delivery and combination therapy across diverse diseases.

## ASSOCIATED CONTENT

### Supporting Information

Materials and Methods, Characterization of synthesized HSA NPs, Size and zeta potential measurements of single drug VAN, SSZ, and EGF-loaded HSA NPs, Size and zeta potential measurements of dual drug (VAN+EGF)- and (SSZ+EGF)-loaded HSA NPs, CD spectra of HSA+SSZ and HSA+VAN at different concentrations of VAN and SSZ, Size and zeta potential measurements of triple drug (VAN+SSZ+EGF)-loaded HSA NPs, the binding free energy data of single-, dual-, and triple-drug loaded HSA NPs.

## Author Contributions

H.C., T.M., and S.Z. designed the research; H.C. and T.M. performed the research; H.C., T.M., A.D., and S.Z. analyzed and interpreted the data; H.C., A.D., T.M., I.O., R.Z., and S.Z. wrote and edited the paper.

### Notes

The authors declare no competing financial interests.

## ACKNOWLEDGEMENTS

This work was supported by the National Institutes of Health (NIH; R01 DK136941 to S.Z.) and the Crohn’s and Colitis Foundation (Litwin IBD Pioneers Award # 993684 to S.Z.). In addition, this work was supported by the UM1TR004408 award (Utilize Harvard University Core Facilities, Award to H.C.) and a KL2/Catalyst Medical Research Investigator Training Award (CMeRIT Award to S.Z.) from Harvard Catalyst | The Harvard Clinical and Translational Science Center (National Center for Advancing Translational Sciences, National Institutes of Health) and financial contributions from Harvard University and its affiliated academic healthcare centers. The content is solely the responsibility of the authors and does not necessarily represent the official views of Harvard Catalyst, Harvard University and its affiliated academic healthcare centers, or the National Institutes of Health. We also thank the Harvard Center for Nanoscale Systems (CNS) for technical support and access to core facilities for nanoscale analysis. We thank Dr. Giovanni Traverso for providing access to laboratory facilities and instrumentation that supported portions of this work. We thank David C. Thorn for assistance with circular dichroism (CD) data collection and helpful discussions related to the study. We thank Bruna Santos for assistance with DLS and HPLC data collection during the early stages of the project. The graphical abstract and the schematics in Figures 1-6 were created with Biorender.

