## Supplementary Material for "Intra-nanoparticle Drug-protein Interactions Mediate Sequential Therapeutic Release"

### Materials and Methods

**Materials.** Human serum albumin (HSA), L-Glutathione reduced (GSH), Vancomycin (VAN, vancomycin hydrochloride from Streptomyces orientalis), Sulfasalazine (SSZ), sodium chloride (NaCl), sodium hydroxide (NaOH), lipopolysaccharide (LPS), ammonium phosphate dibasic, acetonitrile, and ethanol were purchased from Millipore Sigma. Epidermal growth factor (EGF) and EGF ELISA kit were purchased from R&D Systems. UltraPure™ DNase/RNase-Free Distilled Water was purchased from Invitrogen. Dialysis membranes (molecular weight cutoff: 3.5 kDa & MWCO 20 kDa) and Slide-A-Lyzer™ Dialysis Cassettes (20 kDa MWCO) were purchased from Spectrum Laboratories, Inc. and ThermoFisher Scientific, Inc., respectively.

**Synthesis of single drug-loaded HSA NPs.** For SSZ- or EGF-loaded HSA NPs, 225 μL of 80 mg/mL of HAS and 225 μL of 30.8 mg/mL of GSH (both were dissolved in water) were mixed and stirred at 800 rpm under 37°C for 1h. After 1 h incubation, the mixture was allowed to cool to room temperature. For SSZ-loaded NPs, the SSZ solution was prepared at a concentration of 7.4 mg/mL by dissolving 17.5 mg into NaOH (1.75 mL of $10^{-3}$ M NaOH + 0.6 mL of $10^{-1}$ M NaOH). For EGF-loaded NPs, EGF was used at 200 μg/mL dissolved in water. Then, SSZ or EGF drug solutions (50 μL) were added to the mixture and stirred for 10 min. For VAN-loaded NPs, VAN was mixed with GSH solution first at room temperature at a concentration of 2.5 mg/275 uL and 7.7 mg/275 μL, respectively, and mixed with 200 μL HSA solution. To form NPs, 2 mL of 100% ethanol was added dropwise to the mixture. The resulting solutions were then stirred at 1200 rpm for 10 min at 37°C. The resultant NPs were dialyzed against water at 4°C overnight to remove ethanol, GSH, and the unloaded drugs (3.5 kDa MWCO for VAN- and SSZ-loaded NPs and 20 kDa MWCO for EGF-loaded NPs).

**Synthesis of dual drug-loaded HSA NPs.** For (SSZ+EGF)-loaded HSA NPs, similar steps were followed as the single-drug loaded NPs, except for the drug loading. 25 μL of SSZ solution was first added to the mixture of HSA and GSH and stirred for 5 min, followed by the addition of 25 μL EGF solution and further stirring for 5 min. For (VAN+EGF)-loaded HSA NPs, VAN and GSH solutions were first mixed at room temperature at a concentration of 2.5 mg/245 uL and 7.7 mg/245 μL, respectively, and then combined with 180 μL of HSA solution. After incubation for 1h, the mixture was allowed to cool to room temperature. To load EGF, 50 μL EGF solution (100 μg/mL) was added to the mixture and stirred for 10 minutes. To form NPs, 2 mL of 100% ethanol was added dropwise to the mixture. The resulting solutions were stirred at 1200 rpm for 10 min at 37°C. To remove ethanol, the resultant NPs were dialyzed against water at 4°C overnight (MWCO: 3.5 kDa).

**Synthesis of triple drug-loaded HSA NPs.** For (VAN+SSZ+EGF)-loaded HSA NPs, VAN and GSH were first mixed at room temperature at concentrations of 2.5 mg/245 μL and 7.7 mg/245 μL, respectively, and then mixed with 180 μL of HSA solution. The mixture was stirred at 800 rpm in an incubator at 37°C for 1h and subsequently allowed to cool to room temperature. To load SSZ and EGF, 25 μL SSZ solution was added to the mixture and stirred for 5 min, followed by the addition of 25 μL of EGF solution and further stirring for another 5 min. To form NPs, 2 mL of 100% ethanol was added dropwise to the mixture. The resulting mixture was stirred at 1200 rpm for 10 min at 37°C. To remove ethanol, the resultant NPs were dialyzed against water at 4 °C overnight (MWCO: 3.5 kDa).

**Size and zeta potential of NPs**. The hydrodynamic size (Z-average intensity mean) and zeta potential of the NPs were measured using 10 μL of the NPs diluted in 1 mL of 1 mM NaCl. Each sample was measured at least three times at 25℃ using a Malvern Zetasizer Nanoseries. To observe NP morphology, 50 μL of NPs were mixed with 50 μL of 100% ethanol and then added onto a clean silica surface and dried at room temperature with natural convection. The prepared sample was then coated with gold to minimize charging before imaging (Zeiss Gemini 360 FE-SEM).

**Quantification of drug encapsulation**. The drug encapsulation efficiency (EE) of VAN and SSZ was analyzed by HPLC (Agilent 1260 Infinity), while the EE of EGF was quantified by ELISA (R&D Systems). The chromatographic separation was carried out on a 150 mm × 4.6 mm Eclipse XDB-C18 Agilent analytical column with 5 μm spherical particles. EE% = [(the amount of drug in NPs)/(total drug input)] *100%. For SSZ quantification, 500 μL of SSZ-loaded NPs was mixed with 500 μL of 30.8 mL GSH solution in a 1.5 mL microcentrifuge tube, shaking at 300 rpm (VWR® Microplate Shaker) under room temperature overnight. To prepare the samples for HPLC, the mixture was transferred to a 15 mL Falcon® Tube, and then 1.0 mL of 100 mM ammonium phosphate was added and mixed well. Then the mixture was sonicated (Branson 2000) for 5 min and filtered with a 0.2 µm nylon filter (VWR Acrosdic syringe filters). The mobile phase consisted of 70% 100 mM ammonium phosphate and 30% acetonitrile with a flow rate of 1.0 mL/min, an injection volume of 5 µL, a run time of 8 min, a column temperature of 30°C, and UV detection at 254 nm. For VAN quantification, 500 µL of VAN-loaded NPs solution was mixed with 500 µL of 30.8 mg/mL GSH solution at room temperature overnight, shaking at 300 rpm. To prepare the samples for HPLC, the mixture was transferred to a 15 mL Falcon® Tube, and then 1.0 mL of ultrapure water was added and mixed well. Then the mixture was sonicated for 5 min and then filtered with a 0.2 µm nylon filter. The mobile phase consisted of 97% 0.15(v/v) formic acid in water and 3% acetonitrile, with a flow rate of 1.0 mL/min, an injection volume of 10 µL, a run time of 7 min, a column temperature of 30°C, and UV detection at 230 nm. For EGF quantification, 100 µL of EGF-loaded NPs was mixed with 100 µL of 30.8 mg/mL GSH solution in a 1.5 mL microcentrifuge tube at room temperature overnight, shaking at 300 rpm. To prepare the samples for ELISA, 200 µL of ultrapure water was added. EGF concentration was then measured following the manufacturer's ELISA protocol. Absorbance was recorded at 450 nm with a reference wavelength of 570 nm (Tecan Spark).

***In vitro* drug release studies.** To investigate the drug release, supernatant was collected from cultured macrophages with and without LPS treatment. Briefly, RAW 264.7 macrophages were seeded in a 96-well plate (15,000 cells/well) in DMEM containing 10% heat-inactivated fetal bovine serum (HI FBS) at 37°C with 5% $CO_2$. After 24h, 10 µL of 10 ng/mL LPS was added to cells for activation, while 10 µL of medium was added to non-activated cells. Cell culture supernatant was collected after another 24 hours of culture and stored at -20°C for drug release studies. 500 µL single-, dual- and triple-drug loaded NPs and 500 uL media (250 uL MΦ +250 uL water, 100 uL MΦ +400 uL water, 50 uL MΦ +350 uL water, 25 uL MΦ + 475 uL water, and 500 uL water; MΦ: supernatant from LPS-activated macrophages) were mixed. The mixtures were maintained at 37°C under 100 rpm shaking (New Brunswick Innova® 40 Benchtop orbital shaker) throughout the whole experiment. For each time point, samples were centrifuged at 10,000 rpm at 24 °C for 5 min, and the supernatant was analyzed for drug release by HPLC or ELISA, as described above.

**FTIR spectra measurement**. HSA solution with and without drugs (VAN and SSZ) for the single-drug loaded HSA NPs were prepared, and then 100 uL solutions were mixed with pure ethanol. The mixed solution was dropped and naturally dried before FTIR measurement. A Cary 630 FTIR (Agilent Technologies) was used to measure the spectra of HSA and drug-loaded HSA. The spectra (spectral resolution 2 $cm^{-1}$; 60 scans) obtained were converted into transmittance.

**Circular dichroism (CD) measurement.** The CD measurements of pure HSA and HSA mixed with drugs (SSZ and VAN) were performed using a J-1500 spectropolarimeter (Jasco, Tokyo, Japan) over a wavelength range of 180-260 nm. A solution of 5 μM HSA was prepared in water as a control, and 5, 10, 20, and 40 μM of drugs (VAN and SSZ) were added to the HSA solution to study their effect on the HSA chain conformation. CD spectra were recorded at 20°C using a scanning rate of 20 nm/min. All spectra were baseline-corrected by subtracting the corresponding buffer signal.

**Binding free energy measurement using molecular dynamics simulation.** The crystal structure of HSA was obtained from the Protein Data Bank and prepared by removing crystallographic water molecules, co-crystallized ligands, and other heteroatoms. The three-dimensional structures of VAN and SSZ were generated using RDKit and parameterized with the GAFF2 force field using the AM1-BCC charge model. Molecular docking was performed using AutoDock Vina to identify the most favorable binding pose of VAN and SSZ within the HSA binding pocket. The docked complex was subsequently processed in the AMBER ff19SB force field and solvated in an explicit TIP3P water box with 12 Å padding under physiological ionic conditions (0.15 M NaCl). The AMBER topology was converted into GROMACS format for molecular dynamics simulations. Following energy minimization, the system was equilibrated under NVT and NPT ensembles at 300 K and 1 bar using positional restraints before a 100 ns production simulation was carried out without restraints. Trajectory post-processing was performed to remove periodic boundary condition artefacts and recenter the protein-ligand complex. Finally, the binding free energy between VAN/HSA and SSZ/HSA was calculated using the MM-GBSA method.

For the binding energy calculation of the dual-drug loaded NPs, the rigid framework of HSA and EGF served as the receptor for simultaneous docking of VAN and SSZ into their respective, distinct binding pockets. The individual binding free energies for both VAN and SSZ were then calculated via the MM-GBSA method. For the triple-drug loaded HSA NPs, a sequential docking strategy was implemented because the independent docked top-ranked poses of SSZ and VAN substantially overlapped, with a minimum heavy-atom distance of approximately 0.45 Å. To avoid an unrealistic binding configuration, one ligand was fixed within the HSA-EGF complex, and the second ligand was subsequently docked in the presence of the pre-bound ligand. Briefly, SSZ was first docked into the fixed HSA-EGF framework using AutoDock Vina and locked as a rigid component of the receptor matrix. VAN was subsequently docked in the presence of bound SSZ. Alternatively, VAN was first docked into the fixed HSA-EGF framework using AutoDock Vina and locked as a rigid component of the receptor matrix, after which SSZ was subsequently docked in the presence of bound VAN. For both arrangements, the binding free energies of SSZ and VAN were subsequently calculated using the MM-GBSA method.

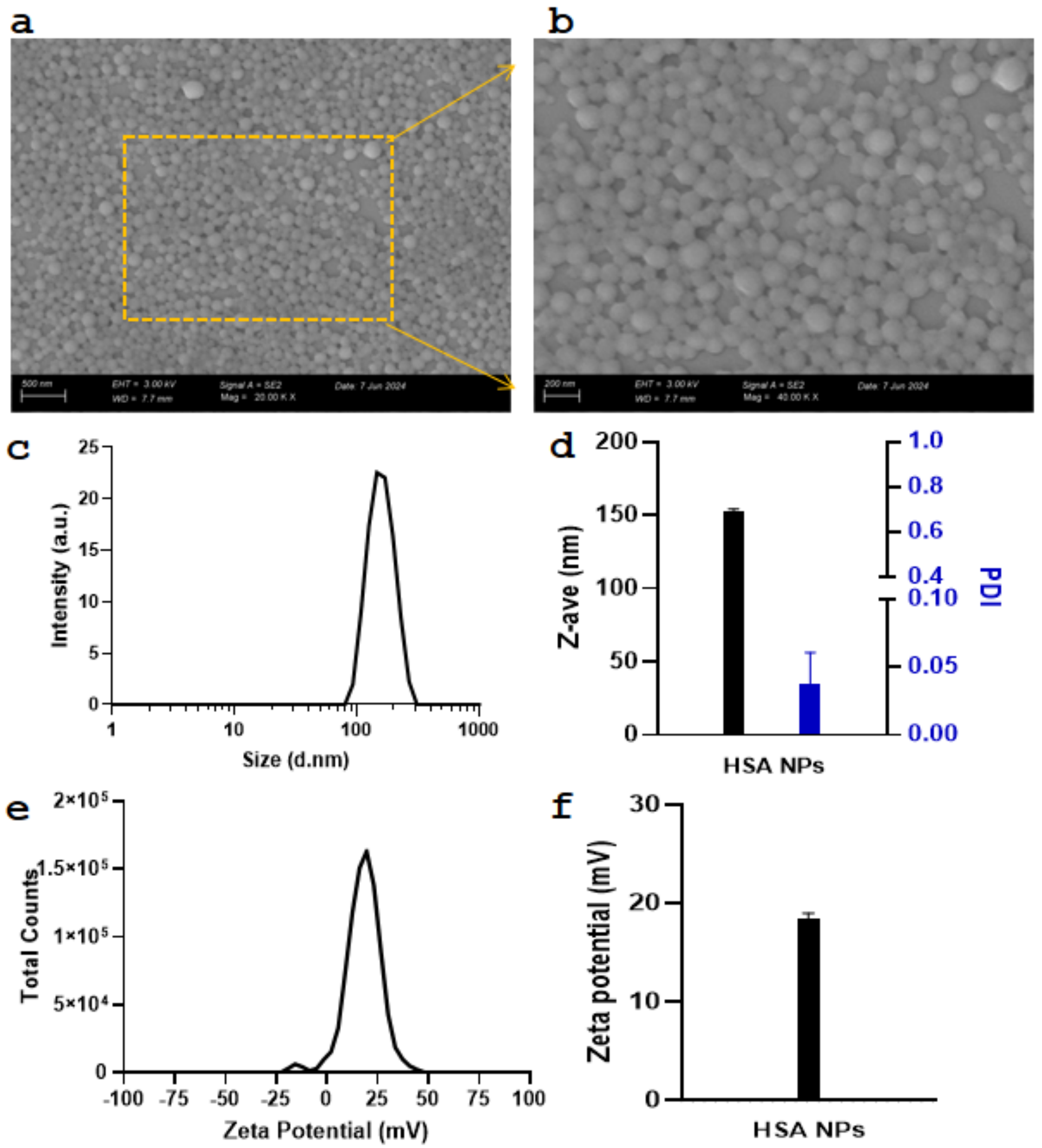


**Figure S1.** Characterization of synthesized HSA NPs. (a, b) Representative SEM images of HSA NPs morphology. (c, d) Hydrodynamic size and PDI of HSA NPs, and (e, f) zeta potential of HSA NPs.

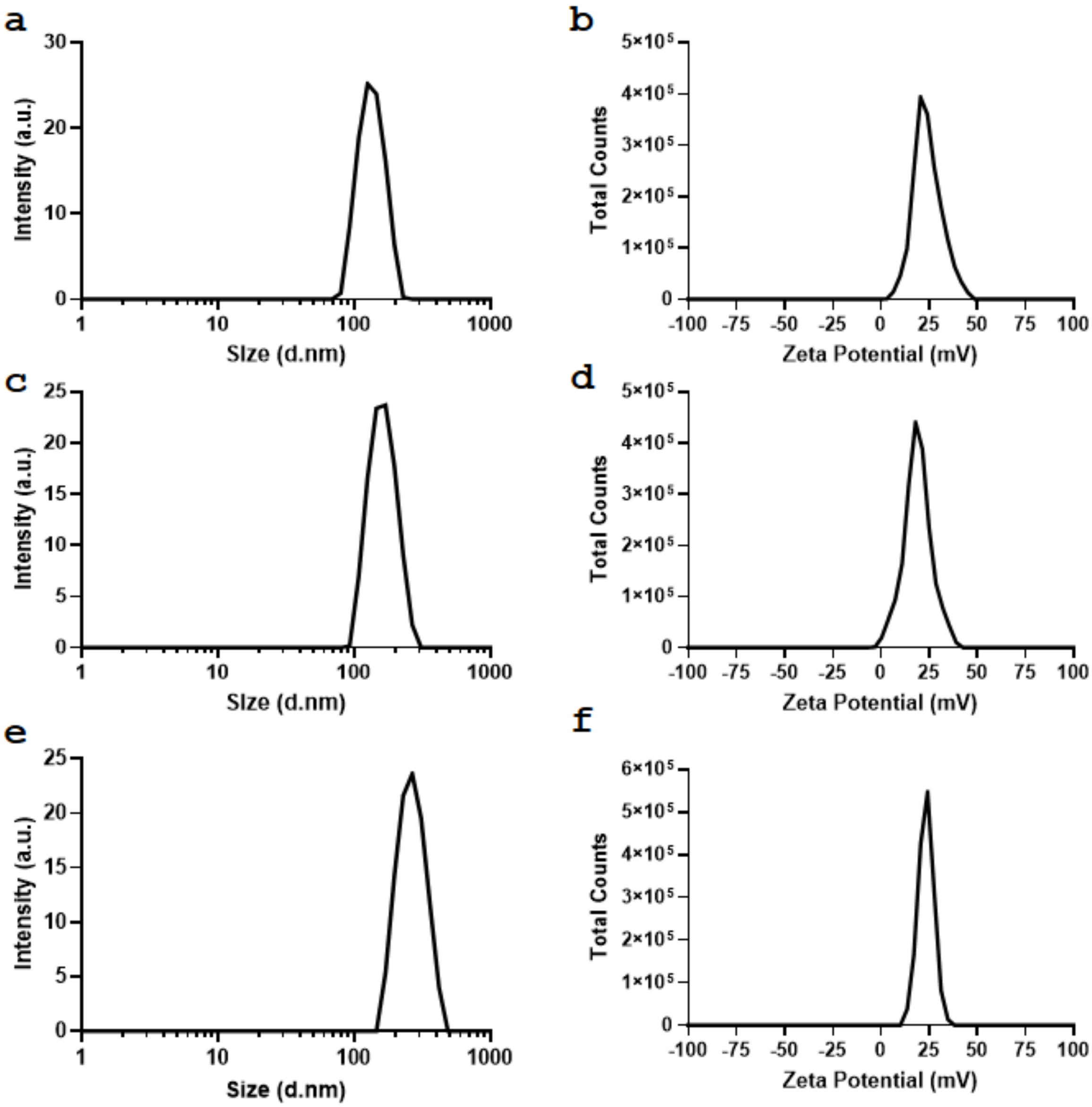


**Figure S2.** Representative size and zeta potential measurements of single drug (a, b) VAN, (c, d) SSZ, and (e, f) EGF-loaded HSA NPs.

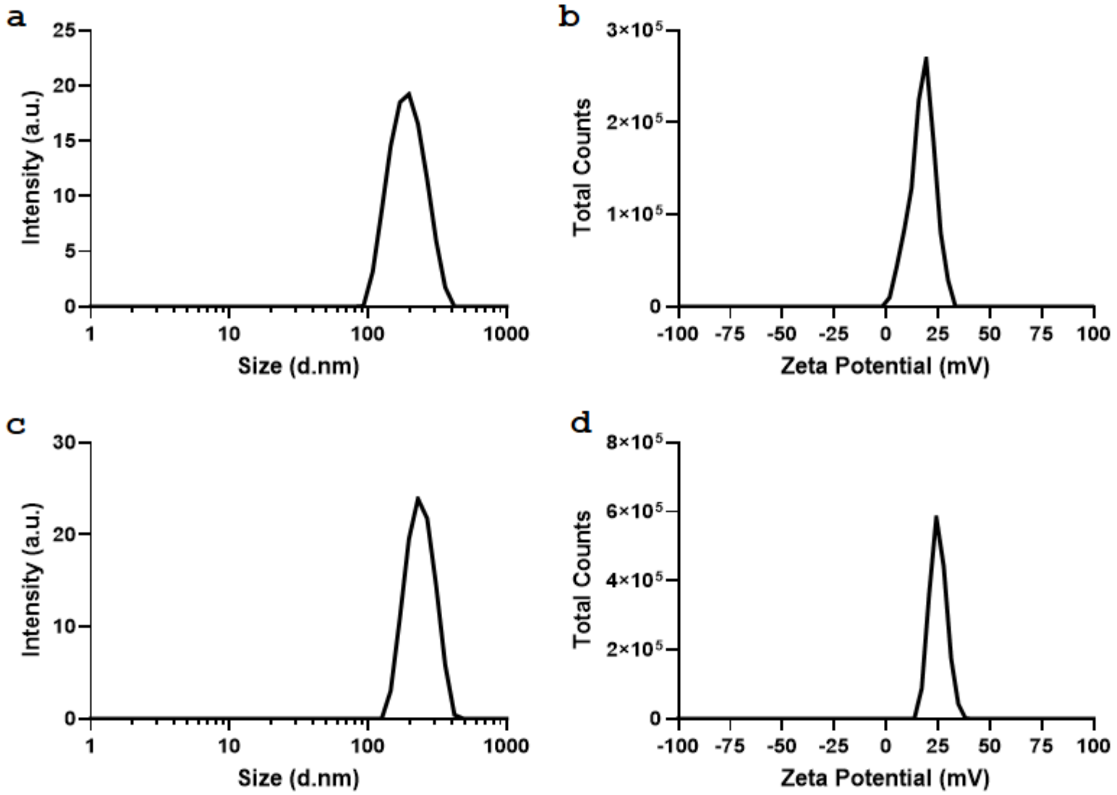


**Figure S3.** Representative size and zeta potential measurements of dual drug (a, b) (VAN+EGF)- and (c, d) (SSZ+EGF)-loaded HSA NPs.

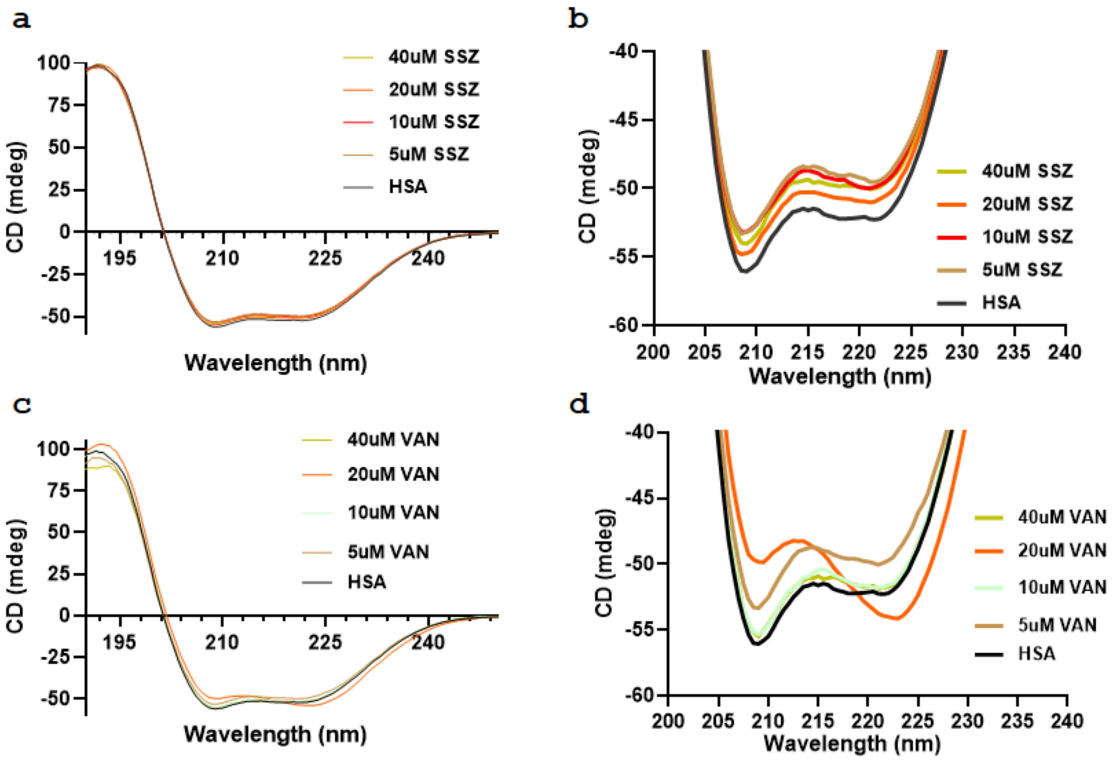


**Figure S4.** Circular Dichroism (CD) Spectra of (a, b) HSA+SSZ and (c, d) HSA+VAN at different concentrations of VAN and SSZ, respectively.

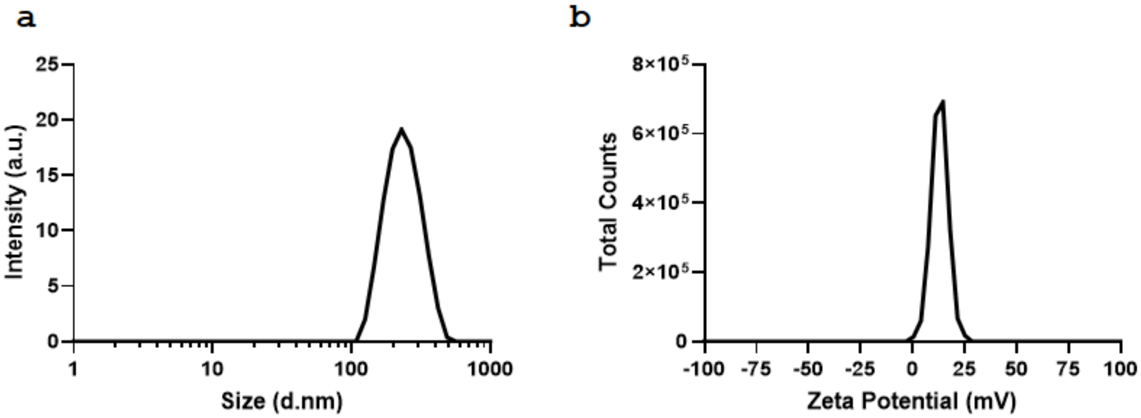


**Figure S5.** Representative (a) size and (b) zeta potential measurements of triple drug (VAN+SSZ+EGF)-loaded HSA NPs.

**Table S1.** The binding free energy data of single-, dual-, and triple-drug loaded HSA NPs. The analysis was performed on a 5001-10001 frame GBSA analysis.

| Loading condition | Ligand evaluated | ΔG GBSA (kcal/mol) | Std. Dev. | Std. Err. |
|---|---|---|---|---|
| Single load | Sulfasalazine | -15.0350 | 6.8816 | 0.3074 |
| Single load | Vancomycin | -23.3051 | 9.0205 | 0.4030 |
| Dual load with EGF | Sulfasalazine | -16.7653 | 5.7733 | 0.2579 |
| Dual load with EGF | Vancomycin | -12.1310 | 7.0540 | 0.3151 |
| Triple load, VAN fixed, evaluating SSZ | Sulfasalazine | -14.5053 | 13.8314 | 0.6179 |
| Triple load, SSZ fixed, evaluating VAN | Vancomycin | -16.6071 | 6.1877 | 0.2764 |
